# Identification and Functional Insights into New Phage Tail-Like Bacteriocins (PTLBs) Targeting *Pseudomonas aeruginosa* as new antimicrobials

**DOI:** 10.1101/2025.08.20.671207

**Authors:** Clara Ibarguren-Quiles, Lucia Blasco, Carla López-Causape, Inés Bleriot, Laura Fernández-García, Lucia Arman, Antonio Barrio-Pujante, Concha Ortiz-Cartagena, Belén Aracil, Luis Mariñas-Pardo, Rafael Cantón, Antonio Oliver, María Tomás

**Author notes:** These researchers contributed equally to the work (Antonio Oliver, María Tomás). **RESEARCH TOPIC(S):** Phage tail-like bacteriocins (PTLBs) as antimicrobial agents.

## Abstract

The current health crisis caused by multidrug resistant (MDR) pathogens is one of the health problems of most concern globally. Infections caused by these pathogens, such as *Pseudomonas aeruginosa*, lead to high rates of complications, particularly in compromised patients such as cystic fibrosis (CF) patients. The need to counteract and minimize the forecast future impact has led to the rescue of phage therapy. The use of bacteriophages has important advantages, including highly specific targeting, self-amplification at the infection site, minimal disruption of the microbiome, safety and biocompatibility. However, the capacity of bacteria to escape these entities results in a form of resistance that compromises the effectiveness of the therapy. This involves the search for potential alternatives, such as the phage tail-like bacteriocins (PTLBs), also named as tailocins. These high molecular weight particles resemble the tail structure of bacteriophages and are characterized by the absence of genetic material, avoiding the development of resistance, one of the major handicaps associated with phage therapy. In this study, we detected 34 different PTLBs in 75 *P. aeruginosa* genomes, with different serotypes and sequence types, 11 of which were characterized as novel F-type PTLBs subtypes (F13-F24). Furthermore, we report that four selected PTLBs (R1, F15, F19, R3-F24) can deal with bacterial infection, with the R1 and the F15 PTLBs being the most efficient in clearing infection *in vitro*, yielding a survival rate of more than 75% in the *Galleria mellonella* larvae *in vivo* model. This reaffirms the potential of PTLBs to control *P. aeruginosa* infections, which can cause chronic infections in some patients, such as people with CF, due to its strong impact as a MDR bacterium.

**GRAPHICAL ABSTRACT:** 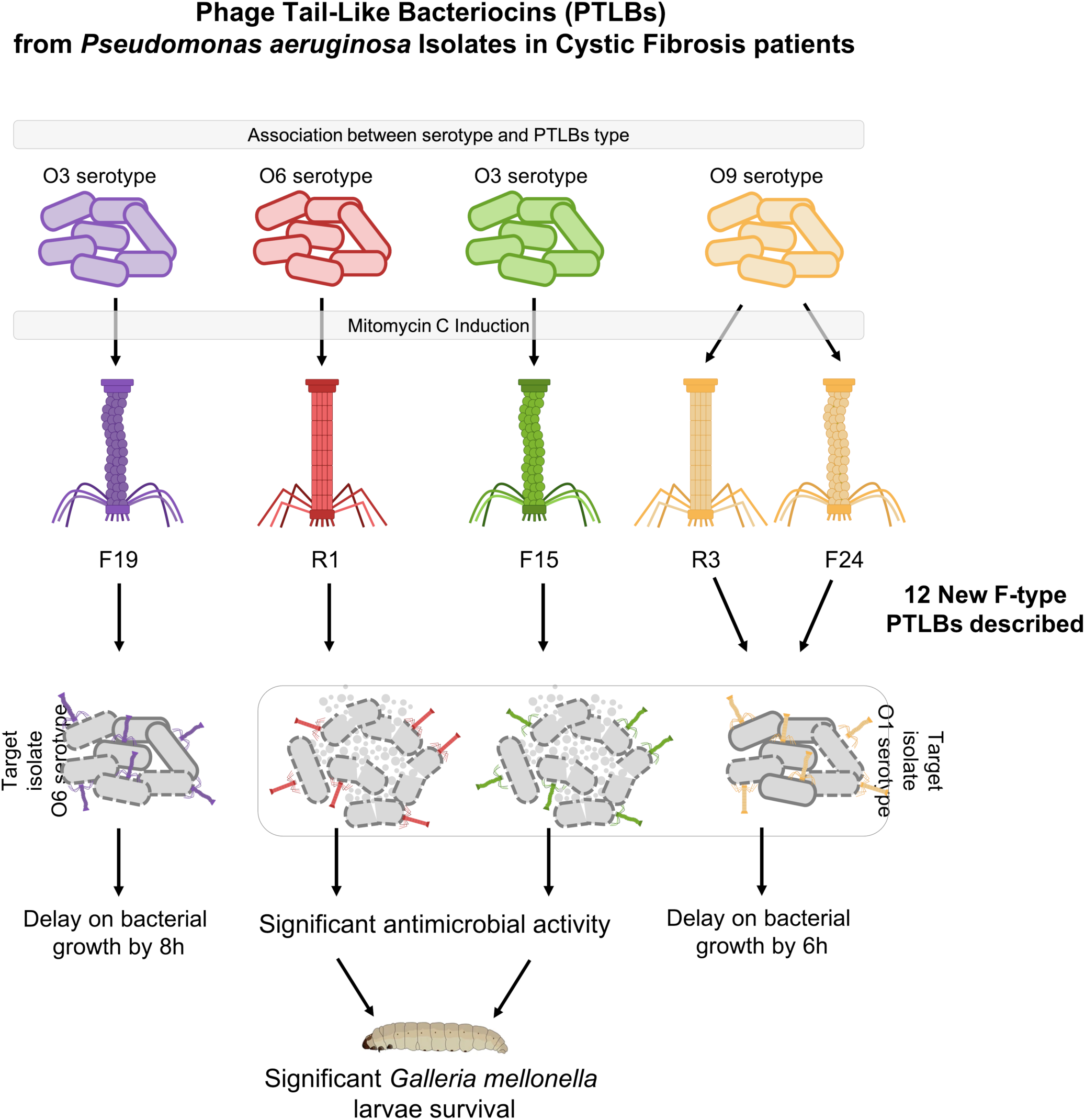

**HIGHLIGHTS:**

-The 75 *Pseudomonas aeruginosa* genomes from people with cystic fibrosis in the study collection included at least one Phage tail-like bacteriocins (PTLB) cluster.
-From the 34 different PTLBs detected in the study collection, 7 were R-type, 10 were complex (R and F-type encoded) and 14 were F-type PTLBs.
-11 new F-type PTLBs were described in the *Pseudomonas aeruginosa* collection under study.
-An association between the O-antigen present on the surface of the *Pseudomonas aeruginosa* isolate and the encoded PTLB subtype was detected.
-The R1 and F15 PTLBs subtypes display high antimicrobial activity both *in vitro* and *in vivo* (*Galleria mellonella*).

## INTRODUCTION

Cystic fibrosis (CF) is an autosomal recessive genetic disease characterized by mutations in the gene coding for a bicarbonate and chloride ion transporter channel known as the CF transmembrane conductance regulator (CFTR) (1, 2). Alterations in the CFTR prevent the immune system from acting efficiently, allowing opportunistic and/or nosocomial pathogens to invade the respiratory tract and cause chronic and recurrent infections (pulmonary involvement is the primary cause of morbidity and mortality in people with CF (pwCF). *Pseudomonas aeruginosa* is one of the pathogens that most frequently appear as an infectious agent of these patients (3, 4). In the 2023 European Cystic Fibrosis Society Patients Registry, it was reported that 19% of pediatric patients and 38% of adult patients had been infected, at least once, by *P. aeruginosa* (5). Currently, the clinically relevant problem associated with these Gram-negative bacterial infections lies in the high incidence of multidrug-resistant (MDR) strains (6, 7).

The lack of efficient antibiotics to combat the MDR infections compromises the health of pwCF, and new treatments are therefore needed (8). Phage therapy, a recently recovered antimicrobial strategy, is based on the antibacterial activity of bacteriophages (viruses that infect bacteria) (9–11). Although several studies have demonstrated the effectiveness of this type of treatment, the development of phage resistance mechanisms is a major drawback (12, 13). Thus, alternative therapeutic approaches such as the use of phage tail-like bacteriocins (PTLBs) are being considered. PTLBs, also known as tailocins (or pyocins when derived from *P. aeruginosa* 14), are high molecular weight protein structures similar to phage tails. They are classified within the large family of bacteriocins, i.e. molecules synthesized and used by bacteria in an environmental bacterial competition, as a survival strategy (15, 16). As a bacterial competitive mechanism, the strategy is activated by the SOS response (17). Once assembled, the PTLBs are released through the activity of endolysins encoded in the lytic cassette, killing the producer cell in an altruistic manner (16, 18). Once released, the PTLBs act by binding to the bacterial O-antigen in the lipopolysaccharide (LPS); the particle then recognizes specific residues within the core region of LPS from the target bacteria, binding and killing the cells by depolarization of the membrane potential, and causing their lysis due to the alteration in cell energetics (19–22) (23, 24).

As indicated by their name, PTLBs resemble the tails of bacteriophages. The absence of capsid and nucleic acid are characteristic and differentiate PTLBs from phages (25), conferring the advantage of reduced development of resistance and lack of horizontal gene transfer. Two types of PTLBs are described depending on whether their structure is rigid /(R-type) or flexible (F-type). Both types of PTLBs are also further subdivided on the basis of genetic characteristics and their spectrum of action. Five subtypes of R-type PTLBs have been described (R1-R5) and twelve subtypes are distinguished by the F-type PTLBs (F1-F12) (26, 27). Thus, bacterial genomic analysis is useful for determining the kind of PTLB that the bacteria can secrete: R-type, F-type or both. In particular, PA01 is used as a reference as it contains both types in its cluster, specifically R2-F2, and, when this occurs, regulatory and lysis genes are shared (28).

Therefore, the main objective of our study was to investigate the presence of these PTLBs in clinical isolates of *P. aeruginosa* from pwCF and to evaluate their potential use to treat bacterial infections, especially those caused by CF *P. aeruginosa*.

## RESULTS

### Genomic identification of PTLBs

Overall 75 *P. aeruginosa* clinical isolates from 25 CF patients were analyzed to search for clusters corresponding to PTLBs. Bioinformatic analysis confirmed the presence of at least one nucleotide sequence corresponding to a PTLB cluster in each of the 75 isolates. A homology study of the sequence identified 34 different PTLBs, each named after its isolate producer. Thereafter, the phylogenetic analysis revealed sequences with high homology, which were clustered in two large groups according to phylogenetic distance (**Figure 1**).

**Figure 1.**
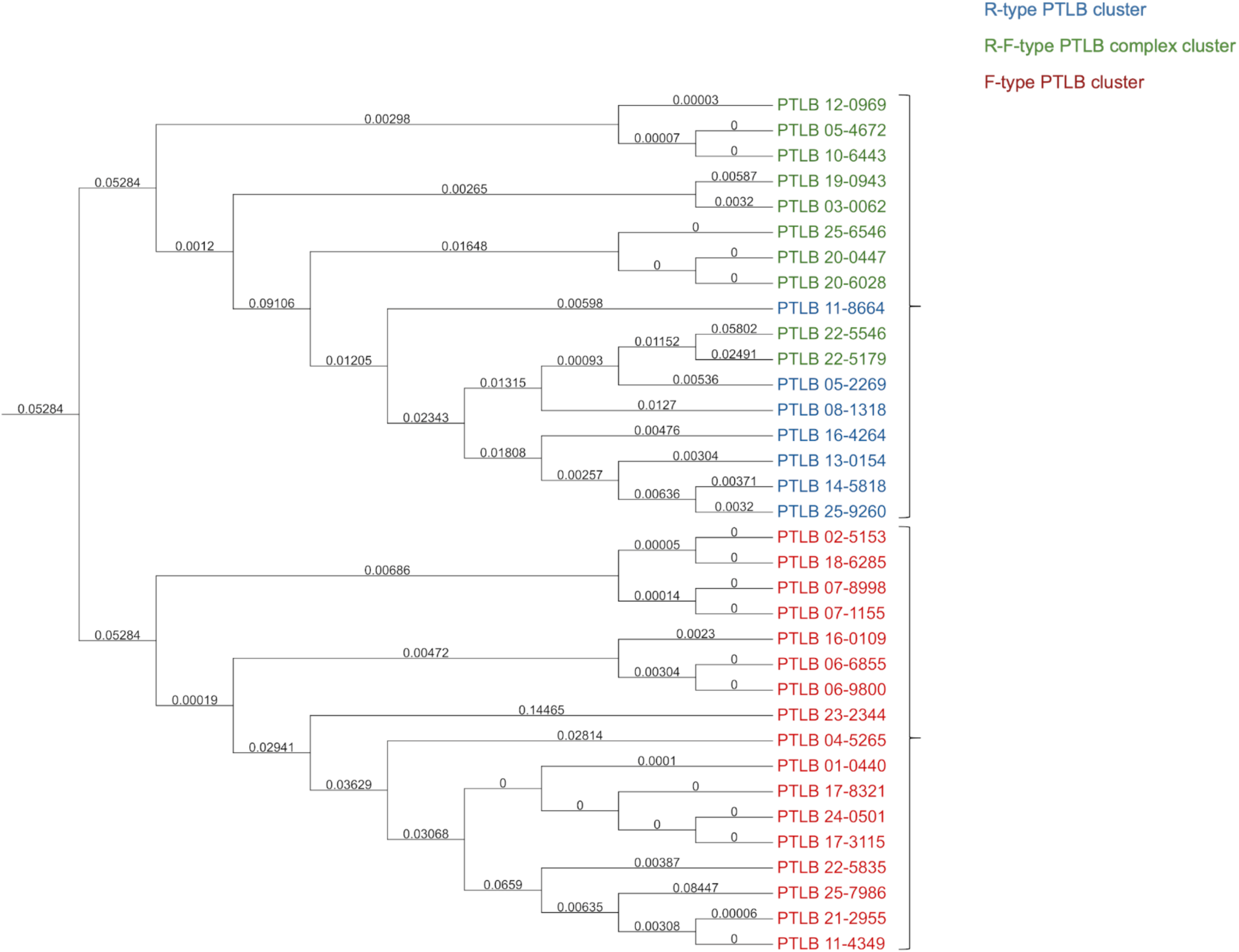
Phylogenetic analysis of 34 distinct PTLBs. The tree shows two different groups: the first is composed of bacterial strains with R-type PTLB clusters (blue) and R-F PTLB complex clusters (green) and the second is composed of bacterial strains with F-type PTLB clusters (red).

### Classification of PTLBs

In order to classify the 34 different PTLBs obtained (**Figure 2**), a proteomic homology study was conducted. The protein organization in the cluster showed whether the cluster encodes typical R-type, F-type proteins or both, here referred to as a PTLB complex. Thus, the collection was found to be composed of 50% of the F-type, 29.41% of PTLB complexes and 20.59% of the R-type. The groups of PTLB types corresponded to the phylogenetic clusters obtained in the phylogenetic tree (**Figure 1**).

**Figure 2.**
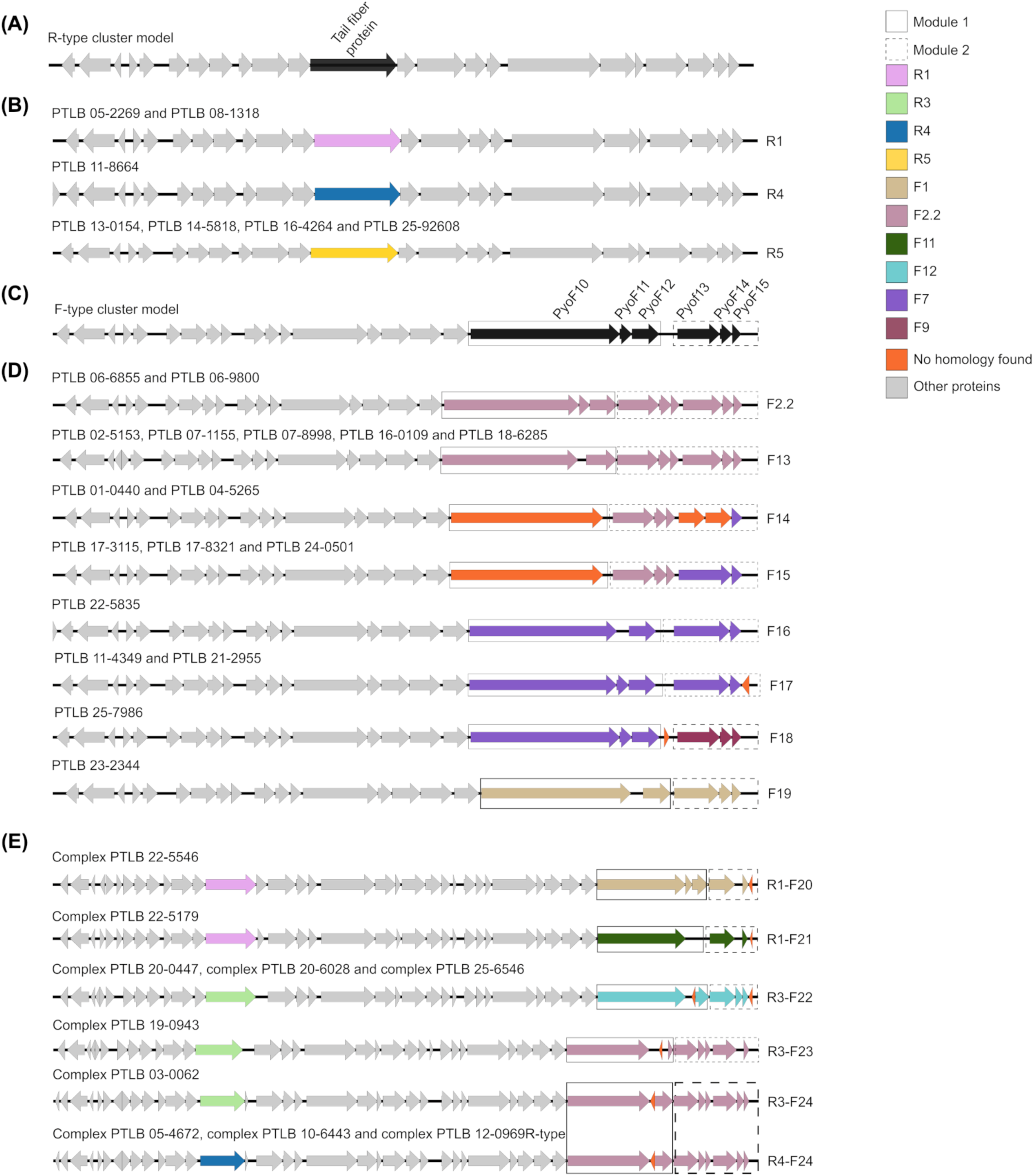
PTLBs subtype classification study. The 34 different PTLB clusters included in this study were subjected to a homology study. The grey arrows correspond to those proteins not used for subtype classification of each of the PTLBs. In addition, the proteins used to determine the type of PTLB is color-coded according to the subtype: R1 (pink), R3 (light green), R4 (navy blue), R5 (maroon), F1 (brown), F2.2 (mauve), F11 (dark green), F12 (turquoise), F7 (purple), F9 (yellow) and unknown homology (orange). The solid black box represents the set of three proteins that comprises module 1. The proteins in module 2 are outlined by a dashed black box.

A homology study was carried out to confirm these results, taking as a reference the tail fibre protein associated with the five subtypes of R-type PTLBs (**Figure 2A**). This enabled identification of two of the R1 subtype, one of the R4 subtype and four of the R5 subtype (**Figure 2B**). In order to classify the F-type PTLBs into subtypes, homology analysis was performed within the last six proteins of the cluster, encoded at the 3’ end (PyoF10, PyoF11, PyoF12, PyoF13, PyoF14 and PyoF15), which are grouped in two modules, as described by Saha et al. (2023) (27) (**Figure 2C)**.The 17 F-type PTLBs identified were grouped into eight different subtypes (**Figure 2D**). The 06-6855 and 06-9800 PTLBs were found to belong to the subtype F2.2 previously identified by Saha et al. (27). The group composed by PTLB 02-5153, PTLB 07-1155, PTLB 07-8998, PTLB 16-0109 and PTLB 18-6285 was classified in the new subtype, F13, which is homologous to the F2.2 subtype but lacks the PyoF11 protein. The other five F-type PTLBs (01-0440, 04-5265, 17-3115, 17-8321 and 24-0501) in the study collection were found to present a characteristic pattern as module 1 only contained the PyoF10 protein, which did not show any homology with any of the PyoF10 proteins previously described. In addition, duplication of the module 2 proteins (Pyo13.2, Pyo14.2 and Pyo15.2), as previously described in the F2.2 subtype, was observed (27). PTLB 01-0440 and PTLB 04-5265 were classified as subtype F14, and two new proteins were found in module 2, in the positions of the Pyo13.2 and Pyo14.2, and a final F7 subtype PyoF15 protein. Regarding PTLB 17-3115, PTLB 17-8321 and PTLB 24-0501, the final two proteins in module 2 were shown to be homologous to the F7 subtype PyoF13 and PyoF15 proteins and were classified as a new PTLBs subtype, F15. Another three F-subtypes showed some homology with the F7 subtype but with some differences that classified them as new subtypes: F16, F17 and F18. Thus, in comparison with the F17 subtype, the protein corresponding to PyoF11 was absent in the F16 subtype (PTLB 22-5835), while in F17 (PTLB 11-4349 and PTLB 21-2955) there was an extra protein at the 3’ end of the module 2. Finally, the F18 subtype (PTLB 25-7986) had an extra protein between the PyoF12 and PyoF13 proteins, and in this PTLB module 2 also showed some homology with the corresponding one present in the F9 subtype. The F-type PTLB 23-2344 presented homology to the F1 subtype and lacked PyoF11 protein.

Moreover, in the present study, ten PTLB complexes were described (**Figure 2E**). An extra protein at the 3’ end of the module 2 was also found in the PTLB complex 22-5546 and the PTLB complex 22-5179. The first was identified as the R1-F20 subtype, with F20 being a new F-subtype with strong homology with the F1 subtype and absence of the PyoF14. The PTLB complex 22-5179 was characterized as the R1-F21 subtype and showed the absence of the proteins PyoF11, PyoF12 and PyoF14, with PyoF10, PyoF13 and PyoF15 homologous to the F11 subtype. The next three PTLB complexes (PTLB complex 20-0447, PTLB complex 20-6028 and PTLB complex 25-6546) comprised proteins homologous to the F12 subtype and a new protein in the PyoF11 position. Additionally, an extra protein was found downstream of PyoF15 in these PTLBs complexes. These characteristic traits resulted in a new subtype of F-subtype PTLB, with these complexes being R3-F22 subtypes. The PTLB 19-0943 complex was classified as R3-F23 by its homology with the R3 subtype and F14 subtype, which was a newly identified subtype characterized by a new protein in the PyoF11 position and a shorter form of the protein PyoF12 and the lack of PyoF14.2. Furthermore, PTLB 03-0062 was characterized as a PTLB R3-F24 subtype. The final three PTLB complexes were identified as the R4-F24 subtype. The F24 subtype characterized as a new subtype showed some homology with the F2.2 subtype but with substitution of the protein PyoF11 for an unknown protein.

### Determination of the PTLB host range

A PTLB host range assay was conducted in order to establish the activity of these particles (**Figure 3**). Thus, one PTLB was selected for each of the subtypes described above for its extraction and subsequent spot test analysis (**Figure 3A**). The host range of the 17 PTLBs over 17 host strains ranged from 0 to 77%. The widest host range corresponded to the R1 PTLBs extracted from O6 serotype isolates (PTLB 05-2269, PTLB 22-5179 and PTLB 22-5546). The PTLB 17-8321 also has a very wide host range (50%), and PTLB 11-8664, PTLB 14-5818 and PTLB 21-2955 did not show any activity on the isolates, while the remaining six had an intermediate host range. Interestingly, the only isolate with serotype O11 was the only one resistant to all of the PTLBs tested.

**Figure 3.**
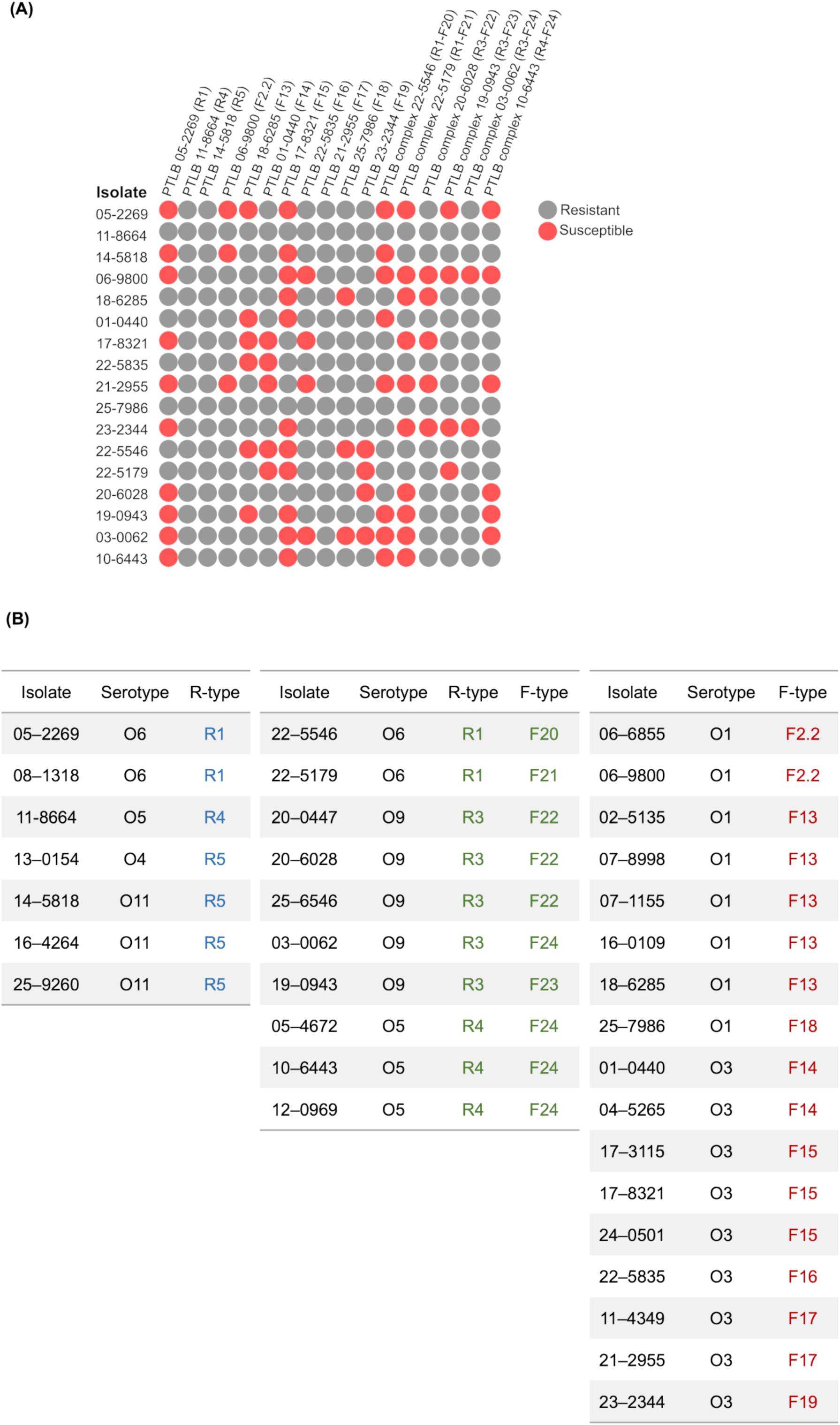
PTLBs host range analysis. (A) Susceptible isolates (red) and resistant isolates (grey) for each PTLB are shown in a 2D histogram constructed using Morpheus (https://software.broadinstitute.org/morpheus). (B) Association between the serotype and encoded PTLB in each of the 34 isolates of study.

A strong correlation between the isolate serotype and its PTLB R subtype was also observed (**Figure 3B**), so that the O6 and O11 serotype isolates carried R1 and R5 PTLB respectively. Regarding the PTLB complexes, the O6 serotype isolates also encoded R1-F, the O9 serotype encoded R3-F and the O5 serotype isolates encoded the R4-F subtypes.

Although this correlation was not observed for the F type PTLBs, this type was shown to be encoded primarily by isolates belonging to the O1 and O3 serotypes.

### Characterization of PTLBs by Transmission Electron Microscopy (TEM)

Based on the results obtained regarding the host range and the PTLBs type, four PTLBs were selected to confirm the R and F-type classification by examination of the morphology by TEM: PTLB 03-0062 (R1-F24), PTLB 23-2344 (F19), PTLB 05-2269 (R1) and PTLB 17-8321 (F15) (**Figure 4**). For PTLB 03-0062 and PTLB 05-2269, uncontracted rigid structures were observed, corresponding to R-type PTLBs, of length 130 and 125 nm and width 20 and 50 nm, respectively. Moreover, some structures appeared in a contracted form in which the core maintained the same length while the sheath reduced its length to 50 nm in both samples (**Figure 4A, 4C**). On the other hand, F-type PTLBs were found to be thinner more flexible structures in PTLB 03-0062, PTLB 23-2344 and PTLB 17-8321, with the first of length 160 nm and width 20 nm and the other two samples of length 120 nm and width 10 nm (**Figure 4A, 4B, 4D**).

**Figure 4.**
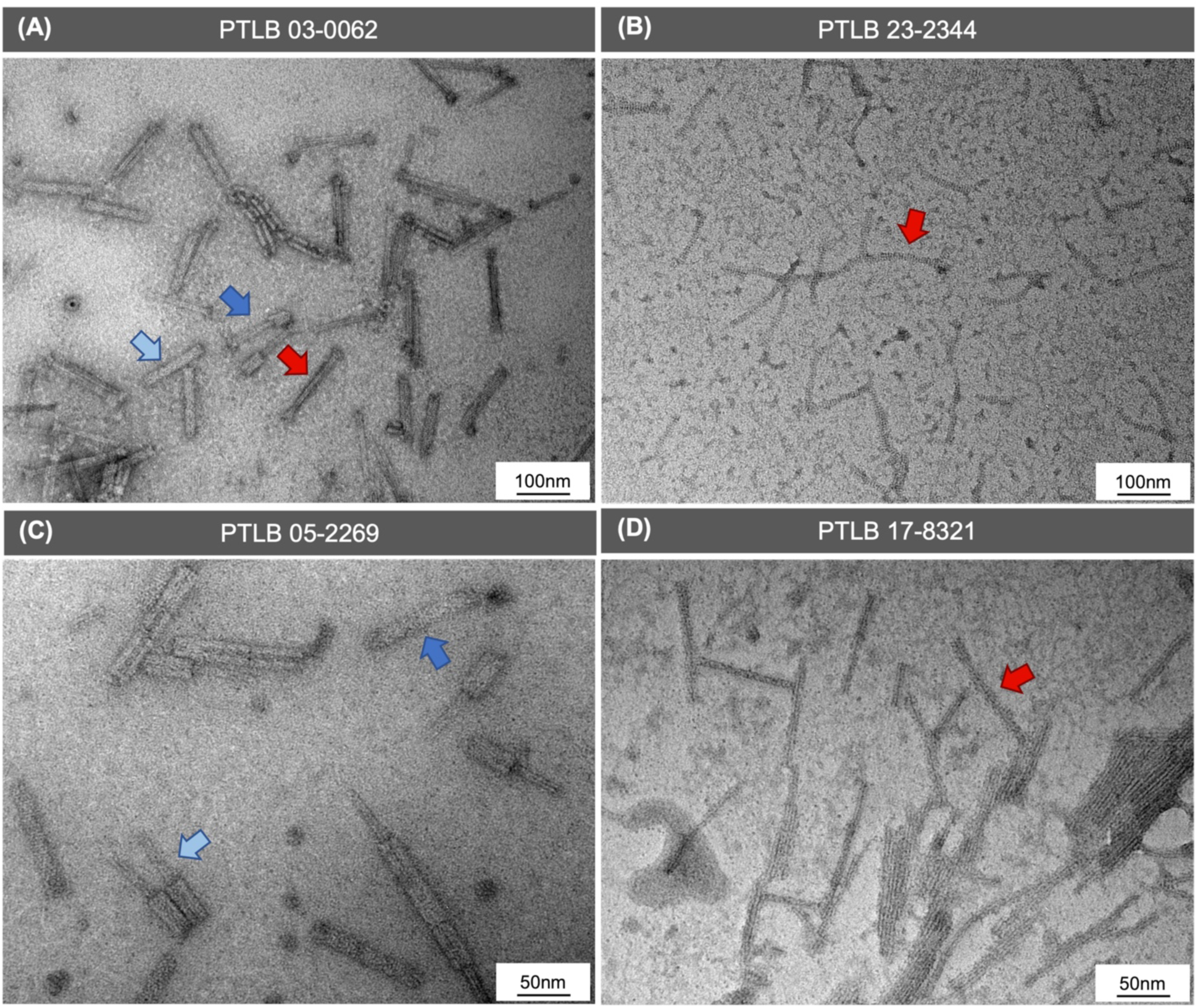
Images representing the PTLBs morphological structure determined by TEM. (A) PTLB complex 03-0062 containing R-type PTLB morphological characteristics in a contracted state (dark blue arrow) and an uncontracted state (light blue arrow) and F-type PTLB morphological characteristics (red arrow). (B) PTLB 23-2344 F-type morphological characteristics (yellow arrow). (C) PTLB 05-2262 R-type morphological characteristics in a contracted state (red arrow) and an uncontracted state (green arrow). (D) PTLB 17-8321 F-type morphological characteristics (yellow arrow).

### Testing the antimicrobial activity of PTLBs

For the purpose of studying the antimicrobial activity of the four previously selected PTLBs, a killing curve assay was performed on a culture of isolates 06-9800 and 22-5546 (**Figure 5**). The results of the killing curves for PTLBs 03-0062 and 23-2344 showed delays in bacterial growth of 6 and 8h, respectively. After 24h incubation, bacterial growth reached the control stationary phase within the PTLB 03-0062 treatment, but not in the case of the PTLB 23-2344, in which there was a significant reduction of 1 log in the CFU counts (**Figures 5A, 5B 5C, 5D**). In PTLB 05-2269 and PTLB 17-8321, the cultures were dead at 6 h with no OD measures and 0 CFU counts, but at 24h, regrowth yielded values 10^3^ and 10^5^ CFU/ml respectively (**Figure 5E, 5F, 5G, 5H**).

**Figure 5.**
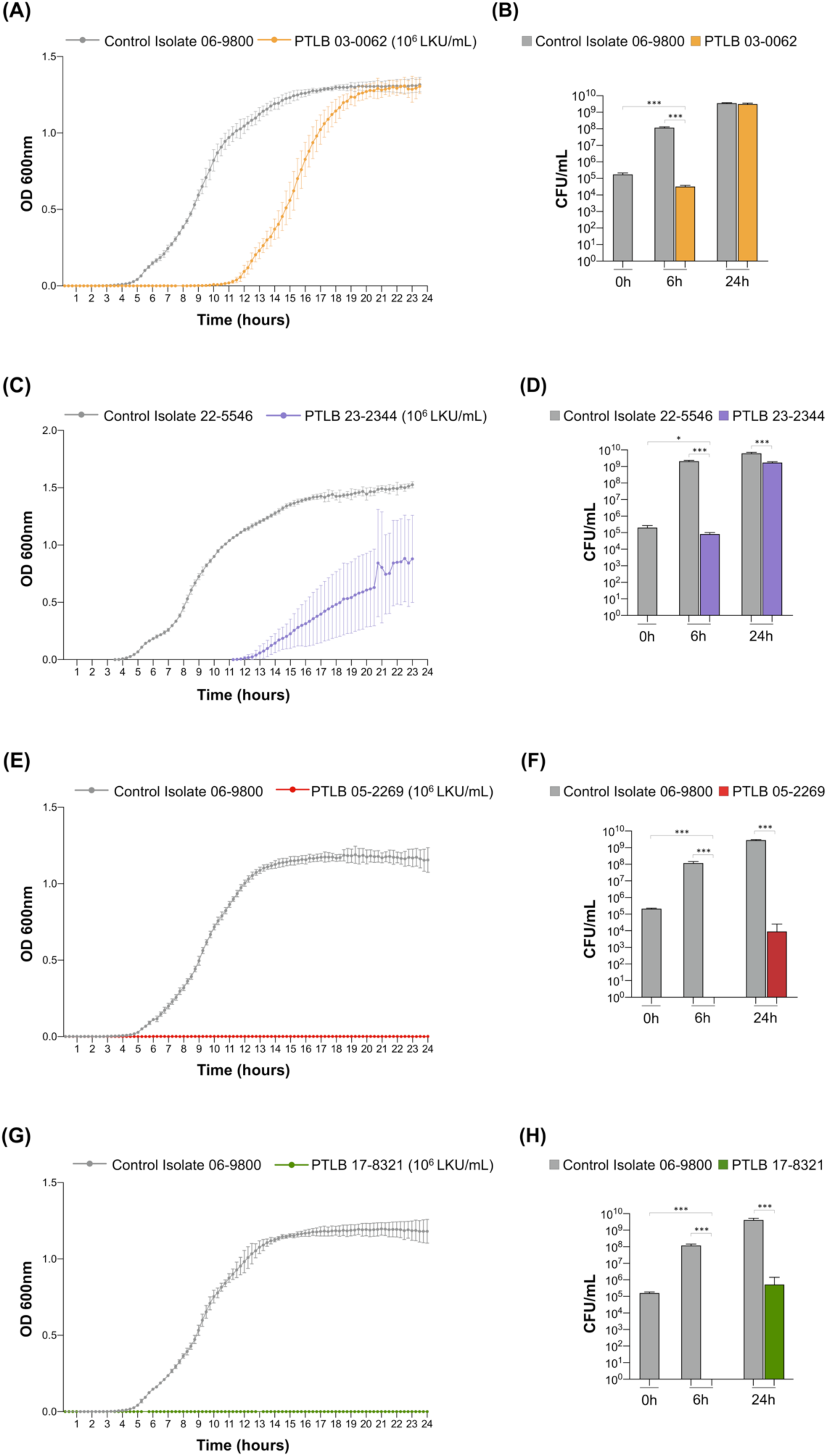
Growth and killing curves and CFU/mL counts. (A) Growth curve of the isolate 06-9800 untreated (grey) and treated with PTLB 03-0062 (orange) during 24 h. (B) CFU/mL count of the isolate 06-9800 untreated (grey) and treated with PTLB 03-0062 (orange) at 0, 6 and 24h. (C) Growth curve of the untreated isolate 22-5546 (grey) and the isolate treated with PTLB 23-2344 (purple) for 24h. (D) CFU/mL count of the untreated 22-5546 isolate (grey) and the isolate treated with PTLB 23-2344 (purple) at 0, 6 and 24 h. (E) Growth curve of the untreated isolate 06-9800 (grey) and the isolate treated with PTLB 05-2269 (red) for 24h. (F) CFU/mL count of the untreated isolate 06-9800 (grey) and the isolate treated with PTLB 05-2269 (red) at 0, 6 and 24 h. (G) Growth curve of the untreated isolate 06-9800 (grey) and the isolate treated with PTLB 17-8321 (green) for 24 h. (H) CFU/mL count of the untreated isolate 06-9800 (grey) and the isolate treated with PTLB 17-8321 (green) at 0, 6 and 24h. Unpaired t-test. (*) P < 0.05. (****) P < 0.0001. Absence of asterisk indicates no statistically significant difference (P > 0.05).

### Antimicrobial activity of PTLBs in the *Galleria mellonella in vivo* model

The antimicrobial activity showed by PTLBs 23-2344, 05-2269 and 17-832, in the killing curves was tested *in vivo* in a *G. mellonella* larvae model (**Figure 6**). Each of the PTLBs were tested in an infection with the host isolate after previous testing to determine the LD_50_. No significant improvement was observed regarding the infection with isolate 22-5546 treated with PTLB 23-2344, although no signs of toxicity were observed in the larvae group inoculated with this PTLB (**Figure 6A**). In survival assays with the other two PTLBs, high survival rates were obtained in the treated larvae, i.e. of 85 % for PTLB 05-2269 and of 90 % PTLB 17-8321 (**Figure 6B; 6C**). Both treatments were also demonstrated to be non-toxic, as observed in the PTLBs survival of the control groups.

**Figure 6.**
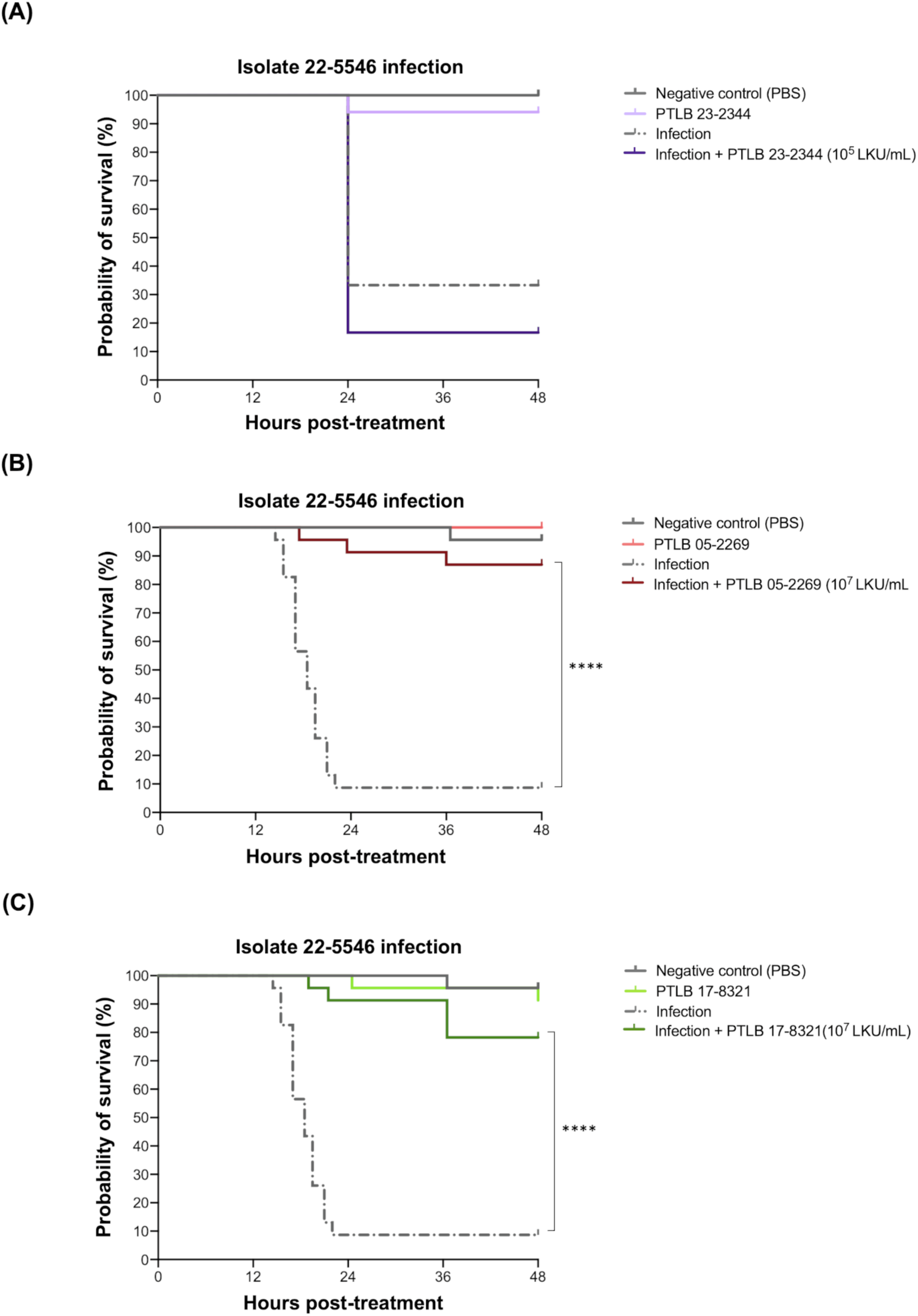
*G. mellonella* survival curves when the infected larvae are untreated or treated with different PTLBs. (A) Percentage survival of *G. mellonella* larvae infected with the isolate 22-5546 alone or treated with the PTLB 23-2344 1h post-infection (B) Percentage survival of *G. mellonella* larvae infected with isolate 06-9800 alone or treated with the PTLB 05-2269 1h post-infection. (C) Percentage survival of *G. mellonella* larvae infected with isolate 06-9800 alone or treated with the PTLB 17-8321 1h post-infection. Mantel-Cox test. (****) for P <0.0001. Absence of asterisk indicates no statistically significant difference (P > 0.05).

## DISCUSSION

PTLBs are competitive weapons that bacteria, such as *P. aeruginosa*, use to compete with other bacteria, commonly different strains from the same species (16, 29). The antimicrobial activity of these protein structures, which are similar to tail phages, can be used in the fight against MDR bacteria. The objectives of the present study were to identify and characterize the activity of different PTLBs against *P. aeruginosa*.

Study of *P. aeruginosa* isolates from pwCF patients detected the presence of at least one PTLB cluster in each of the isolates, suggesting the importance of these protein structures in the CF competitive environment. This confirms the findings of a previous study conducted by our research group in which a similar prevalence of PTLBs sequences was observed in other *P. aeruginosa* collection (21). Annotation of the PTLBs genome and the homology study revealed 34 PTLBs with the characteristic structure, with regulatory, structural and lysis cassettes (30, 31), and in this case three were encoded in clusters for the R type, eight for the F type and six combined a R type and a F type in the cluster with a shared regulatory cassette, comprised between the genes *trpD* and *trpE* (28) This classification was confirmed by a rigid or flexible morphology observed by TEM.

The phylogenetic tree revealed two main groups, one consisting of PTLB complexes and R-type PTLBs and the other consisting of the F-type PTLBs. The clustering of the R type and the complexes is due to the high homology of the R subtypes; thus, the PTLB 11-8664 (R4) branch was found after the three PTLB complexes R4-F and before the two PTLB complexes R1-F (22-5546 and 22-5179), which, in turn, are located in a branch together with PTLB 05-2269 and PTLB 08-1318, both of the R1 subtype.

The R and F type PTLBs are subdivided into several subtypes differentiated by some specific genes (25).The following subtypes of the R-type PTLBs were identified: R1, R3, R4 and R5, previously characterized by the tail fiber protein (32). The F-subtypes were characterized on the basis of the C-terminal region, which is variable and enables distinction of each subtype (27). Analysis of the F subtypes identified 12 new subtypes. Five PTLBs had similarities in module 1, with the main differences being in the PyoF11 protein, which was substituted by a hypothetical protein with a reverse gene reading pattern in the F23 and F24 subtypes. This protein was absent in F13, F14, F15, F16, F19 and F21. Classification of the F-subtype showed that module 1 was more variable regarding the number of proteins than the module 2, in which the PyoF14 protein was absent in F16, F17, F20 and F21 subtypes. The PyoF14.2 protein was also absent in the F21 and F23 subtypes. Although the PyoF11 and PyoF12 proteins, present in module 1, have been considered essential for the functionality and specificity of F-type PTLBs (27), we noted that their absence did not affect their activity or specificity. Thus, the PTLBs lacking this protein, such as PTLB 17-8321, showed a wide host range and a significant bactericidal activity.

Additionally, differences in host range between PTLB 03-0062 (R3 and PTLB 10-6443, both of which share the F4 type PTLB, but differ in the R type [R3 or R4]), indicate the importance of the R type PTLB in recognition of the target strain. Furthermore, the R1 subtype, either alone or accompanied by an F-type, yielded a similar pattern. Although the R5 subtype has been reported to display the widest host range, PTLB 17-8321 (F15), PTLB 05-2265 (R1 subtype) and the PTLB 22-5179 complex (R1-F20) showed the widest host range (19).

In addition, in a study based on a collection of *P. aeruginosa* isolates from pwCF the populations were quite susceptible to R2 subtype PTLBs (33). Thus, the work conducted by Mei et al. (2023) (33) also demonstrated the high potential of R2 subtype PTLBs as antimicrobial treatments, because irrespective of the level of diversity in the LPS, these isolates were still susceptible to R2. These findings led to the idea that as well as the presence of specific antigens, the physical composition and arrangement of LPS are key factors in PTLBs activity (34). The PTLB target is recognized by its receptor binding proteins (RBPs) which can consist of tail tips, tail fibers or tail spikes, which specifically target the O antigen in the LPS moiety (34). The O antigen type (serotype) and the PTLBs subtype produced, which also implies the acquisition of resistance to this PTLBs from the strains with the same serotype (19). The correlation between PTLB and the serotype producer strain was observed for the R type PTLBs identified, including the complex PTLBs. A weaker correlation was observed for the F type. The correlation between the PTLB type and the serotype and therefore the host range is due to the genetic variability in the R and F subtypes and the host range (16). Blasco et al. (2023) also reported the relationship between the serotype and the PTLB complexes. The serotype of an isolate is thus a determining factor in the putative bactericidal action of a PTLB. Our findings are also consistent with those of a previous study in which isolates with O6 serotype encode for R1 subtypes and are, in turn, resistant to the action of R1 subtypes PTLBs (19). In all cases, the isolates were resistant to the encoded PTLBs, except for PTLB 05-2269, a previously reported exception which may be related to the killing mechanism of PTLBs. The Phage tail-like bacteriocins (PTLBs) depolarize the outer membrane, then puncture and inject the tail structure, in a mechanism related to temporal variations in the LPS density and the spontaneous ability of the PTLs to pass through the O antigen and contract close to the cell envelope (34, 35). Regarding the isolates with O1 and O3 serotypes, a tendency for the PTLB clusters to be absent was reported (19). However, in our collection eight isolates with F-type PTLB clusters were encoded by O1 isolates and the other nine F-type PTLB clusters were detected in the bacterial chromosome of O3 isolates. Thus, the data provided shows a direct link between serotype and F-type PTLB, which has not previously been observed because of the relatively few studies on this flexible structures.

PTLBs show great potential as antimicrobial agents that could be used as alternatives to antibiotics in the fight of AMR. This potential was tested in four of the PTLBs identified in this work, which together covered all the serotypes present in this *P. aeruginosa* collection.

Strong antimicrobial activity was observed *in vitro* for three of the PTLBs tested with 0 counts at 6 h and a significantly low regrowth at 24h. The reduced resistance obtained for PTLBs in comparison with phages, which are also potential antimicrobial agents, is probably due to the absence of genetic material, the main driver of the anti-phage defense mechanisms, thereby reducing the resistance to PTLBs in mutations in the genes encoding the O antigen (34). The results obtained in the killing curves were confirmed in the *in vivo* model of *G. mellonella* larvae. Strong antimicrobial activity was observed for two PTLBs, 17-8321 and 05-2269, as the survival of the treated larvae was significantly higher than the group of infected larvae, reaching as high as 80%. These results, together with the absence of toxicity in this model, indicate PTLBs to be strong candidates for the treatment of infections caused by *P. aeruginosa*. PTLBs have not yet been tested in patients, as opposed to phage therapy (36, 37). Nevertheless, *in vivo* mouse models have shown that natural R2-type PTLBs have a strong capacity to control bacterial infections, supporting the findings of pre-clinical studies (26, 38). Our study findings further reinforce these findings by providing data on how an R5 subtype PTLB and an F24 subtype PTLB improve the survival of *G. mellonella* larvae infected by *P. aeruginosa*, thus demonstrating the therapeutic potential of PTLBs of clinical origin.

## METHODS AND MATERIALS

### Bacterial isolates and culture conditions

A total of 75 clinical isolates of *P. aeruginosa* obtained from 25 CF patients and belonging to 26 Sequence Types (STs) were included in the study (**Table 1**). The isolates were provided by the research group led by Dr Antonio Oliver at the Son Espases University Hospital, Palma de Mallorca (Spain). In an earlier study, the genomes of this collection were analyzed by Next-generation sequencing (NGS), with the MiSeq sequencing system (Illumina platform), and assembled using the Newbler Roche assembler and Velvet (Velvet v1.2.101) (39, 40). The serogroups of the isolates were determined on the basis of the O-specific antigen (OSA) gene cluster sequences, by using PAst 1.0 (41) (**Table 1**).

**Table 1.** Characterization of isolates and Genebank annotation. The O-antigen and the ST of the three isolates of each of the 25 CF patients were determined.

| Patient | *P. aeruginosa* CF isolate | O-antigen | ST |
| --- | --- | --- | --- |
| 01 | 01-0440 | O3 | 1089 |
|  | 01-5978 | O3 | 1089 |
|  | 01-7071 | O3 | 1089 |
| 02 | 02-5135 | O1 | 312 |
|  | 02-5867 | O1 | 312 |
|  | 02-6433 | O1 | 312 |
| 03 | 03-0062 | O9 | 285 |
|  | 03-5302 | O9 | 285 |
|  | 03-5453 | O9 | 285 |
| 04 | 04-5265 | O3 | 274 |
|  | 04-7991 | O3 | 274 |
|  | 04-8869 | O3 | 274 |
| 05 | 05-2269 | O6 | NEW1 |
|  | 05-2840 | O6 | NEW1 |
|  | 05-4672 | O5 | 360 |
| 06 | 06-6855 | O1 | 242 |
|  | 06-7209 | O1 | 242 |
|  | 06-9800 | O1 | 242 |
| 07 | 07-1155 | O1 | 279 |
|  | 07-5966 | O1 | 279 |
|  | 07-8998 | O1 | 279 |
| 08 | 08-1318 | O6 | NEW2 |
|  | 08-4371 | O6 | NEW2 |
|  | 08-5924 | O6 | NEW2 |
| 09 | 09-3048 | O6 | 1109 |
|  | 09-0786 | O6 | 1109 |
|  | 09-9593 | O6 | 1109 |
| 10 | 10-6443 | O5 | 360 |
|  | 10-6518 | O5 | 360 |
|  | 10-7858 | O2 | 360 |
| 11 | 11-4349 | O3 | 198 |
|  | 11-7257 | O5 | 2475 |
|  | 11-8664 | O5 | 2475 |
| 12 | 12-0969 | O5 | 277 |
|  | 12-2742 | O5 | 277 |
|  | 12-2760 | O5 | 277 |
| 13 | 13-0154 | O4 | 1123 |
|  | 13-1387 | O4 | 1123 |
|  | 13-2748 | O4 | 1123 |
| 14 | 14-4114 | O11 | 319 |
|  | 14-4688 | O11 | 319 |
|  | 14-5818 | O11 | 319 |
| 15 | 15-4963 | O6 | 412 |
|  | 15-7676 | O6 | 412 |
|  | 15-8860 | O6 | 412 |
| 16 | 16-0109 | O1 | 252 |
|  | 16-2856 | O1 | 252 |
|  | 16-4264 | O11 | 408 |
| 17 | 17-0755 | O3 | 274 |
|  | 17-3115 | O3 | 274 |
|  | 17-8321 | O3 | 274 |
| 18 | 18-6285 | O1 | 312 |
|  | 18-7126 | O1 | 312 |
|  | 18-9439 | O1 | 312 |
| 19 | 19-0943 | O9 | 1092 |
|  | 19-6618 | O9 | 1092 |
|  | 19-9746 | O9 | 1092 |
| 20 | 20-0447 | O9 | 1072 |
|  | 20-3695 | O9 | 1072 |
|  | 20-6028 | O9 | 1072 |
| 21 | 21-2955 | O3 | 198 |
|  | 21-4234 | O3 | 198 |
|  | 21-9889 | O3 | 198 |
| 22 | 22-5179 | O6 | NEW3 |
|  | 22-5546 | O6 | 2101 |
|  | 22-5835 | O3 | 1134 |
| 23 | 23-2344 | O3 | 701 |
|  | 23-6966 | O3 | CC701 |
|  | 23-9557 | O3 | 701 |
| 24 | 24-0501 | O3 | 274 |
|  | 24-1092 | O3 | 274 |
|  | 24-7416 | O3 | 274 |
| 25 | 25-6546 | O9 | 1072 |
|  | 25-7986 | O1 | 235 |
|  | 25-9260 | O11 | 1613 |

The *P. aeruginosa* isolates were grown in Luria-Bertani (LB) broth medium (0.5% yeast extract; 0.5% NaCl; 1% tryptone) at 37 °C and 180 rpm. When necessary, the LB medium was supplemented with 1.5% agar to produce solid medium. The PTLBs were further characterized by the double agar method, which consists of a base with TA medium (1% tryptone, 0.5% NaCl, 1.5% agar) below a suspension of soft TA agar (1% tryptone, 0.5% NaCl, 0.4% agar).

### Identification and annotation of PTLB clusters *in silico*

The genomic sequences of the 75 CF isolates of *P. aeruginosa* were annotated by RAST (42). A search for the term “pyocin” was used to locate the contig with the PTLB. The *trpD* and *trpE* flanking genes were then located and used to identify the cluster ends (16). Once identified, the PTLBs sequences were also annotated using HMMER (version v3.4) (43). The GenBank accession numbers are the following: PV920217 (PTLB 11-8664), PV940334 (PTLB 05-2269), PV940335 (complex PTLB 22-5546), PV940336 (complex PTLB 22-5179), PV940337 (complex PTLB 20-6028), PV940338 (complex PTLB 19-0943), PV940339 (complex PTLB 03-0062), PV940340 (complex PTLB 10-6443), PV940341 (PTLB 06-9800), PV940342 (PTLB 18-6285), PV940343 (PTLB 01-0440), PV940344 (PTLB 17-8321), PV940345 (PTLB 22-5835), PV940346 (PTLB 21-2955), PV940347 (PTLB 25-7986), PV940348 (PTLB 23-2344) and PV976849 (PTLB 14-5818).

### PTLBs homology and phylogenetic analysis

The homology of the different PTLBs found in each of the 75 isolates was determined using the Nucleotide Blast tool (44). The genomic sequences from the PTLBs identified were also phylogenetically compared by alignment, with the default options in Clustal Omega in the EMBL-EBI server (45). The sequences were then used to construct a Maximum-Likelihood (ML) phylogenetic tree using the Viral Genome Tree Service tool with the default parameters in BV-BRC (version 3.49.1) (46). The tree output was edited using the Interactive Tree Of Life (iTOL) bioinformatic tool (47).

### PTLBs classification into type and subtype

The amino acid sequence corresponding to tail fiber protein (i.e. the component of the PTLB determining its host specificity) of the different R-subtypes were used for classification: R1 (ARI05994.1), R2 (AAG04009.1), R3 (ABP93392.1), R4 (ABP93394.1), R5 (ABP93396.1). (48). The R2-subtype corresponds to the PA0620 protein in the reference strain PA01. In addition, the reference sequences of the six proteins, located in the C-terminal of the cluster, were used to classify the F-type PTLBs subtypes. The six proteins that characterize the F-subtypes were then grouped in two modules: PyoF10, PyoF11 and PyoF12, the proteins belonging to module 1, and PyoF13, PyoF14 and PyoF15, the proteins belonging to module 2, as previously described by Saha et al. (2023) (27).

### Extraction, concentration and purification of PTLBs

The extraction, concentration and purification of each of the PTLBs detected was performed using an adaptation of a previous method (21). Briefly, an overnight culture of the selected isolates was diluted (1:100) and incubated at 37°C and 180 rpm. Once the cultures reached an optical density (OD_600nm_) of 0.3-0.4, 10 µg/ml of mitomycin C (Sigma-Aldrich) was added and the cultures were incubated until they cleared. To ensure maximum bacterial lysis, the lysed cultures were treated with 1% chloroform for 20 minutes at room temperature and centrifuged at 3,700 rpm for 10 min. The supernatants containing the PTLBs were filtered (0.45 µm, FILTER-LAB®PES Syringe filter). The supernatants containing the PTLBs were concentrated by precipitation overnight at 4°C with 10% PEG (polyethylene glycol 6000; Sigma-Aldrich) and 0.5 M NaCl. On the following day, the PTLBs were collected by centrifugation for 15 min at 11,000 rpm and 4 °C. The supernatants were discarded, and the pellets were resuspended in SM buffer (0.1 M NaCl, 10 mM MgSO4, 20 mM Tris–HCl, pH 7.5). An equal volume of chloroform was then added to the final volume (1:1) of each sample, and the mixture was then incubated with gentle shaking for 20 min, before being centrifuged for 10 min at 4,000 rpm. The aqueous phases were recovered and incubated with 10 μg/mL proteinase K at 56 °C for 1 hour, in order to degrade putative prophages induced by mitomycin C. Finally, the PTLBs were purified and dialyzed with a 100 KDa Amicon® Ultra-15 (Merck Milipore) system. The PTLB solutions were stored at 4 °C until use.

### Transmission electron microscopy (TEM) imaging of PTLBs

PTLBs containing solution were examined by TEM (JEOL JEM-1011) after fixing the solution in a grid and negatively stained in 1% aqueous uranyl acetate for 5 min.

### PTLB killing spectrum – host range assay

The host range of thirteen PTLBs was determined in a spot test (49), by exposing them to the same thirteen isolates from the *P. aeruginosa* collection from which the PTLBs were extracted, method. The double agar method was used, in which the upper layer contains the *P. aeruginosa* test isolate, previously grown (at 37°C and shaking conditions) until reaching an OD_600nm_ of 0.6 Abs, mixed with TA soft medium and poured over solid TA medium. Once the top layer had solidified, 3 µL drops containing the purified, diluted PTLB suspension (1:10, 1:1000, 1:100000) were deposited on the surface. Each PTLB killing spectrum was performed in triplicate. A result was considered positive when a clear spot was obtained after incubating the plates overnight at 37 °C.

### Quantification of PTLB killing

The killing activity of PTLBs was indirectly quantified by determining the lethal killing units per milliliter (LKU/mL) using Poisson’s distribution, as previously described (50–53). First, an overnight culture of the target bacteria was diluted 1:100 and grown at 37°C and 180 rpm until reaching an OD_600nm_ of 0.2, corresponding to a concentration of 10^8^ CFU/mL. Once this condition was reached, 100 μL aliquots of serial dilutions (1, 1:10, 1:1000; 1:10000) of the PTLB suspension were mixed with 900 μL of the target bacteria and incubated for 40 min at 37°C. As a control group, the target bacteria were mixed with SM buffer. Each condition was then serially diluted and plated on LB agar and incubated at 37°C until the next day when the number of CFUs were counted. The following formula was used to establish the LKU/mL:

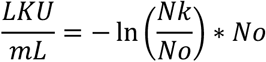

where N_k_ represents the lowest dilution of the PTLB suspension that shows killing activity (with a low number of CFUs) and N_o_ represents the lowest dilution of the PTLB suspension that shows no killing activity (the value is similar to that obtained in the negative control).

### Time killing curve assay exposing bacteria to PTLBs

For the selected PTLB-host combinations that yielded positive spot tests results, killing curve analysis was carried out in LB broth medium. For this purpose, an overnight culture of the selected *P. aeruginosa* clinical host isolates was diluted 1:100 in liquid LB medium and incubated at 37°C at 180 rpm, until reaching an OD_600nm_ of 0.2 Abs, which corresponds to 10^8^ CFU/mL of the host isolate. At this point, a 96-well flat-bottom plate was prepared with 200 μL per well containing 10^5^ CFU/mL, mixed with 10^6^ LKU/mL of the PTLB suspension. In addition, a growth control was tested with the isolate alone. The plates were incubated with continuous orbital shaking at 37°C for 24h hours and read at 600 nm wavelength every 15 min, in a microplate reader (EPOCH, BioTek, Winooski, VT, USA). The bacterial concentration was also determined, at 0, 6 and 24h, by analysis of the CFU/mL at each time. Triplicate samples of both conditions were collected and serially diluted in PBS and subsequently plated, in duplicate, on LB agar plates for subsequent CFU quantification. The data obtained were analyzed using the Unpaired T-test, with the default parameters, and the graphs were plotted using GraphPad Prism Version 8.4.3 (471).

### PTLB toxicity and efficacy of treatment determined by an *in vivo Galleria mellonella* assay

The PTLB toxicity and treatment efficacy were assessed using larvae of the greater wax moth *G. mellonella*, as an *in vivo* model, based on previously described protocols (54–56). The larvae were obtained from a local pet shop (Harkito, Madrid). The 50% lethal dose (LD_50_), which represents the concentration of inoculum that causes death of 50% of the larvae 24 h post-infection, was first determined by infecting several groups of larvae with 10 µL of different dilutions of inoculum. For preparation of the bacterial inoculum, 4 mL of LB broth medium was incubated overnight at 37°C with shaking at 180 rpm. The culture was then centrifuged (4000 x g for 15 min) and washed three times with 4 mL of PBS. Groups of 18 unspotted and healthy *G. mellonella* larvae (n=18), were then infected with 10 µL of the LD_50_ by injecting in the last left proleg, with a Hamilton syringe (Hamilton, Shanghai, China).

After one hour of infection, the groups of infected larvae were treated with 10^3^ LKUs/mL of PTLBs. In addition, one group of infected larvae were injected with PBS to establish a control infection group. The PBS was injected into the last right proleg of each larva. The toxicity associated with the PTLB treatment was evaluated by injecting 10 µL of the mixture into several groups of larvae inoculated one hour earlier with PBS, instead of a bacterial infection. Finally, a control group injected with PBS only, simulating infection and treatment, was included to verify that the injection alone did not affect the larval state. All groups of larvae were placed in Petri dishes and incubated in darkness at 37°C. The number of larvae without motor response to touch (i.e. dead larvae) was recorded regularly over a period of 48 h. Survival curves were plotted using GraphPad Prism Version 8.4.3 (471) and compared using the Log-rank (Mantel-Cox) test.

## ACKNOWLEDGEMENTS

We are grateful to GAIN for the funding provided. L. Fernandez-Garcia and A. Barrio-Pujante were supported by Xunta de Galicia Postdoctoral Grant IN606C-2024/004 and by Xunta de Galicia Predoctoral Grant IN606A-2023/017.

## AUTHOR CONTRIBUTIONS

C.I. conducted the experiments and wrote the original manuscript; L.B., C.L-C., I.B., collaborated in conducting the experiments; L.F-G., L.A., A.B-P. examined the cited studies and collaborated in writing the manuscript; L.M-P., R.C., A.O., revised the manuscript; and finally. M.T. supervised all results, validated the work and obtained funding for the research.

## DECLARATIONS

We declare that there are no conflicts of interest.

## REFERENCES

1. Elborn JS. Cystic fibrosis. Lancet. 2016;388(10059):2519–31.

2. Rey MM, Bonk MP, Hadjiliadis D. Cystic Fibrosis: Emerging Understanding and Therapies. Annu Rev Med. 2019;70:197–210.

3. Gilligan PH. Microbiology of airway disease in patients with cystic fibrosis. Clin Microbiol Rev. 1991;4(1):35–51.

4. Malhotra S, Hayes D, Wozniak DJ. Cystic Fibrosis and *Pseudomonas aeruginosa*: the Host-Microbe Interface. Clin Microbiol Rev. 2019;32(3).

5. Zolin A AA, Bakkeheim E, et al. ECFSPR Annual Highlights Report 2023. 2025.

6. Colque CA, Albarracín Orio AG, Feliziani S, Marvig RL, Tobares AR, Johansen HK, et al. Hypermutator *Pseudomonas aeruginosa* Exploits Multiple Genetic Pathways To Develop Multidrug Resistance during Long-Term Infections in the Airways of Cystic Fibrosis Patients. Antimicrob Agents Chemother. 2020;64(5).

7. Daruka L, Czikkely MS, Szili P, Farkas Z, Balogh D, Grézal G, et al. ESKAPE pathogens rapidly develop resistance against antibiotics in development *in vitro*. Nat Microbiol. 2025;10(2):313–31.

8. Levin-Reisman I, Brauner A, Ronin I, Balaban NQ. Epistasis between antibiotic tolerance, persistence, and resistance mutations. Proc Natl Acad Sci U S A. 2019;116(29):14734–9.

9. Bleriot I, Pacios O, Blasco L, Fernandez-Garcia L, Lopez M, Ortiz-Cartagena C, et al. Improving phage therapy by evasion of phage resistance mechanisms. JAC Antimicrob Resist. 2024;6(1):dlae017.

10. Lin DM, Koskella B, Lin HC. Phage therapy: An alternative to antibiotics in the age of multi-drug resistance. World J Gastrointest Pharmacol Ther. 2017;8(3):162–73.

11. Chan BK, Stanley G, Modak M, Koff JL, Turner PE. Bacteriophage therapy for infections in CF. Pediatr Pulmonol. 2021;56 Suppl 1:S4–S9.

12. Loc-Carrillo C, Abedon ST. Pros and cons of phage therapy. Bacteriophage. 2011;1(2):111–4.

13. Labrie SJ, Samson JE, Moineau S. Bacteriophage resistance mechanisms. Nat Rev Microbiol. 2010;8(5):317–27.

14. Jacob F. Induced biosynthesis and mode of action of a pyocine, antibiotic produced by *Pseudomonas aeruginosa*. Ann Inst Pasteur (Paris). 1954;86(2):149–60.

15. Granato ET, Meiller-Legrand TA, Foster KR. The Evolution and Ecology of Bacterial Warfare. Curr Biol. 2019;29(11):R521–R37.

16. Scholl D. Phage Tail-Like Bacteriocins. Annu Rev Virol. 2017;4(1):453–67.

17. Penterman J, Singh PK, Walker GC. Biological cost of pyocin production during the SOS response in *Pseudomonas aeruginosa*. J Bacteriol. 2014;196(18):3351–9.

18. Vacheron J, Heiman CM, Keel C. Live cell dynamics of production, explosive release and killing activity of phage tail-like weapons for *Pseudomonas* kin exclusion. Commun Biol. 2021;4(1):87.

19. Köhler T, Donner V, van Delden C. Lipopolysaccharide as shield and receptor for R-pyocin-mediated killing in *Pseudomonas aeruginosa*. J Bacteriol. 2010;192(7):1921–8.

20. Kocíncová D, Lam JS. A deletion in the wapB promoter in many serotypes of *Pseudomonas aeruginosa* accounts for the lack of a terminal glucose residue in the core oligosaccharide and resistance to killing by R3-pyocin. Mol Microbiol. 2013;89(3):464–78.

21. Blasco L, de Aledo MG, Ortiz-Cartagena C, Blériot I, Pacios O, López M, et al. Study of 32 new phage tail-like bacteriocins (pyocins) from a clinical collection of *Pseudomonas aeruginosa* and of their potential use as typing markers and antimicrobial agents. Sci Rep. 2023;13(1):117.

22. Mei M, Estrada I, Diggle SP, Goldberg JB. R-pyocins as targeted antimicrobials against *Pseudomonas aeruginosa*. NPJ Antimicrob Resist. 2025;3(1):17.

23. Backman T, Burbano HA, Karasov TL. Tradeoffs and constraints on the evolution of tailocins. Trends Microbiol. 2024;32(11):1084–95.

24. Uratani Y, Hoshino T. Pyocin R1 inhibits active transport in *Pseudomonas aeruginosa* and depolarizes membrane potential. J Bacteriol. 1984;157(2):632–6.

25. Ibarguren C, Bleriot I, Blasco L, Fernández-García L, Ortiz-Cartagena C, Arman L, et al. The world of phage tail-like bacteriocins: State of the art and biotechnological perspectives. Microbiol Res. 2025;295:128121.

26. Scholl D, Martin DW. Antibacterial efficacy of R-type pyocins towards *Pseudomonas aeruginosa* in a murine peritonitis model. Antimicrob Agents Chemother. 2008;52(5):1647–52.

27. Saha S, Ojobor CD, Li ASC, Mackinnon E, North OI, Bondy-Denomy J, et al. F-Type Pyocins Are Diverse Noncontractile Phage Tail-Like Weapons for Killing. J Bacteriol. 2023;205(6):e0002923.

28. Nakayama K, Takashima K, Ishihara H, Shinomiya T, Kageyama M, Kanaya S, et al. The R-type pyocin of *Pseudomonas aeruginosa* is related to P2 phage, and the F-type is related to lambda phage. Mol Microbiol. 2000;38(2):213–31.

29. Backman T, Latorre SM, Symeonidi E, Muszyński A, Bleak E, Eads L, et al. A phage tail-like bacteriocin suppresses competitors in metapopulations of pathogenic bacteria. Science. 2024;384(6701):eado0713.

30. Shinomiya T, Shiga S, Kageyama M. Genetic determinant of pyocin R2 in *Pseudomonas aeruginosa* PAO. I. Localization of the pyocin R2 gene cluster between the *trpCD* and *trpE* genes. Mol Gen Genet. 1983;189(3):375–81.

31. Matsui H, Sano Y, Ishihara H, Shinomiya T. Regulation of pyocin genes in *Pseudomonas aeruginosa* by positive (prtN) and negative (prtR) regulatory genes. J Bacteriol. 1993;175(5):1257–63.

32. Michel-Briand Y, Baysse C. The pyocins of *Pseudomonas aeruginosa*. Biochimie. 2002;84(5-6):499–510.

33. Mei M, Pheng P, Kurzeja-Edwards D, Diggle SP. High prevalence of lipopolysaccharide mutants and R2-pyocin susceptible variants in *Pseudomonas aeruginosa* populations sourced from cystic fibrosis lung infections. Microbiol Spectr. 2023;11(6):e0177323.

34. Carim S, Azadeh AL, Kazakov AE, Price MN, Walian PJ, Lui LM, et al. Systematic discovery of *pseudomonad* genetic factors involved in sensitivity to tailocins. ISME J. 2021;15(8):2289–305.

35. Mei M, Thomas J, Diggle SP. Heterogenous Susceptibility to R-Pyocins in Populations of *Pseudomonas aeruginosa* Sourced from Cystic Fibrosis Lungs. mBio. 2021;12(3).

36. Onallah H, Hazan R, Nir-Paz R, Brownstein MJ, Fackler JR, Horne B, et al. Refractory *Pseudomonas aeruginosa* infections treated with phage PASA16: A compassionate use case series. Med. 2023;4(9):600–11.e4.

37. Jault P, Leclerc T, Jennes S, Pirnay JP, Que YA, Resch G, et al. Efficacy and tolerability of a cocktail of bacteriophages to treat burn wounds infected by *Pseudomonas aeruginosa* (PhagoBurn): a randomised, controlled, double-blind phase 1/2 trial. Lancet Infect Dis. 2019;19(1):35–45.

38. Merrikin DJ, Terry CS. Use of pyocin 78-C2 in the treatment of *Pseudomonas aeruginosa* infection in mice. Appl Microbiol. 1972;23(1):164–5.

39. Blasco L, Ibarguren-Quiles C, López-Causape C, Armán L, Barrio-Pujante A, Bleriot I, et al. Study of the probability of resistance to phage infection in a collection of clinical isolates of *Pseudomonas aeruginosa* in relation to the presence of Pf phages. Microbiol Spectr. 2025;13(3):e0301024.

40. Sastre-Femenia M, Fernández-Muñoz A, Gomis-Font MA, Taltavull B, López-Causapé C, Arca-Suárez J, et al. antibiotic susceptibility profiles, genomic epidemiology and resistance mechanisms: a nation-wide five-year time lapse analysis. Lancet Reg Health Eur. 2023;34:100736.

41. Thrane SW, Taylor VL, Lund O, Lam JS, Jelsbak L. Application of Whole-Genome Sequencing Data for O-Specific Antigen Analysis and In Silico Serotyping of *Pseudomonas aeruginosa* Isolates. J Clin Microbiol. 2016;54(7):1782–8.

42. Overbeek R, Olson R, Pusch GD, Olsen GJ, Davis JJ, Disz T, et al. The SEED and the Rapid Annotation of microbial genomes using Subsystems Technology (RAST). Nucleic Acids Res. 2014;42(Database issue):D206–14.

43. Potter SC, Luciani A, Eddy SR, Park Y, Lopez R, Finn RD. HMMER web server: 2018 update. Nucleic Acids Res. 2018;46(W1):W200–W4.

44. Altschul SF, Gish W, Miller W, Myers EW, Lipman DJ. Basic local alignment search tool. J Mol Biol. 1990;215(3):403–10.

45. Madeira F, Madhusoodanan N, Lee J, Eusebi A, Niewielska A, Tivey ARN, et al. The EMBL-EBI Job Dispatcher sequence analysis tools framework in 2024. Nucleic Acids Res. 2024;52(W1):W521–W5.

46. Olson RD, Assaf R, Brettin T, Conrad N, Cucinell C, Davis JJ, et al. Introducing the Bacterial and Viral Bioinformatics Resource Center (BV-BRC): a resource combining PATRIC, IRD and ViPR. Nucleic Acids Res. 2023;51(D1):D678–D89.

47. Letunic I, Bork P. Interactive Tree of Life (iTOL) v6: recent updates to the phylogenetic tree display and annotation tool. Nucleic Acids Res. 2024;52(W1):W78–W82.

48. Ghequire MG, De Mot R. Ribosomally encoded antibacterial proteins and peptides from *Pseudomonas*. FEMS Microbiol Rev. 2014;38(4):523–68.

49. Kutter E. Phage host range and efficiency of plating. Methods Mol Biol. 2009;501:141–9.

50. Yao GW, Duarte I, Le TT, Carmody L, LiPuma JJ, Young R, et al. A Broad-Host-Range Tailocin from *Burkholderia cenocepacia*. Appl Environ Microbiol. 2017;83(10).

51. Williams SR, Gebhart D, Martin DW, Scholl D. Retargeting R-type pyocins to generate novel bactericidal protein complexes. Appl Environ Microbiol. 2008;74(12):3868–76.

52. Baltrus DA, Clark M, Hockett KL, Mollico M, Smith C, Weaver S. Prophylactic Application of Tailocins Prevents Infection by. Phytopathology. 2022;112(3):561–6.

53. Kageyama M, Ikeda K, Egami F. Studies of a pyocin. III. Biological properties of the pyocin. J Biochem. 1964;55:59–64.

54. Bleriot I, Blasco L, Fernández-Grela P, Fernández-García L, Armán L, Ibarguren C, et al. Studies *in vitro* and *in vivo* of phage therapy medical products Targeting Clinical Strains of *K.pneumoniae* belonging to the clone ST512. Antimicrob Agents Chemother. 2025:e0193524.

55. Peleg AY, Jara S, Monga D, Eliopoulos GM, Moellering RC, Mylonakis E. *Galleria mellonella* as a model system to study *Acinetobacter baumannii* pathogenesis and therapeutics. Antimicrob Agents Chemother. 2009;53(6):2605–9.

56. Blasco L, Tomás M. Use of *Galleria mellonella* as an Animal Model for Studying the Antimicrobial Activity of Bacteriophages with Potential Use in Phage Therapy. Methods Mol Biol. 2024;2734:171–80.

